# Investigation of Protein Melting Temperature Prediction with Cross-Method Validation on Biophysical Data

**DOI:** 10.64898/2026.05.07.723192

**Authors:** Karen Pailozian, Pavel Kohout, Jiri Damborsky, Stanislav Mazurenko

## Abstract

**Motivation:** Protein melting temperature (*T*_*m*_) prediction accelerates the discovery of thermostable enzymes which are crucial for industrial biotechnology often requiring harsh reaction conditions. Experimental determination of *T*_*m*_ remains labour-intensive and varies across techniques, motivating the development of *in silico* predictors. Mass-spectrometry datasets such as Meltome Atlas now enable large-scale *T*_*m*_ prediction with models based on deep learning, but model generalisation across diverse experimental datasets has not been systematically tested.

**Results:** We evaluated the generalisability of state-of-the-art deep learning approaches and explored ESM-based embeddings for *T*_*m*_ prediction. To this end, we assembled the ProMelt training dataset (45 441 proteins) and five independent biophysics-based validation datasets. Our analysis revealed substantial differences between proteomics- and biophysics-based *T*_*m*_ measurements, highlighting the challenge of cross-domain generalisation. Existing state-of-the-art predictors trained on large-scale proteomics datasets showed reduced performance on biophysics-based validation sets. Our fine-tuned embedding-based models, particularly LoRA-adapted ESM-2 (TmProt 1.0), outperformed state-of-the-art predictors in identifying thermostable proteins (*T*_*m*_ *≥* 60 °C) across heterogeneous datasets, achieving AUC scores of 0.75–0.77. We also demonstrated that the available models could be used efficiently in the sequence prioritization task.

**Availability:** The TmProt web server is available at https://loschmidt.chemi.muni.cz/tmprot/. Source code and data are available at https://github.com/loschmidt/TmProt.

## 1 Introduction

Environmental conditions directly influence protein functioning and folding. As many biotechnological processes operate under extreme pH and temperature conditions, discovering biocatalysts with enhanced stability beyond the native environment is often essential (Erdem and Woodley, 2024; Hon et al., 2020). Moreover, protein engineering campaigns targeting other properties, for example, catalytic rates, often lead to mutations compromising protein stability (Hou et al., 2023), necessitating the availability of a stable template. Thus, tools for predicting melting temperatures (*T*_*m*_) of proteins from sequences can accelerate the identification of thermostable proteins that meet the reaction prerequisites (Musil et al., 2019).

Several experimental techniques exist to determine protein *T*_*m*_, each relying on different principles. Biophysical techniques such as differential scanning calorimetry (DSC) (Johnson, 2013), differential scanning fluorimetry (DSF) (Gao et al., 2020), and circular dichroism (CD) (Ranjbar and Gill, 2009) spectroscopy have long been used as a gold standard for evaluating protein stability in wet lab experiments. They typically probe the thermodynamic and structural properties of proteins to determine their *T*_*m*_ by changing the temperature and registering changes in the signal triggered by protein denaturation. High sample consumption and demanding data analysis protocols limit the throughput of such methods to up to several dozen proteins measured in a single protein characterisation campaign (Vasina et al., 2022).

In contrast, proteomics-based methods are higher-throughput, enabling thermostability measurements for thousands of proteins in parallel. For example, thermal proteome profiling (TPP) (Mateus et al., 2016; Jarzab et al., 2020) and limited proteolysis coupled with mass spectrometry (LiP-MS) (Leuenberger et al., 2017) are proteomics-based methods that utilise mass spectrometry to measure protein stability at a large scale. The recently released Meltome Atlas (Jarzab et al., 2020) employed TPP to collect ~48 000 data points across multiple organisms, the scope well beyond the reach of classical biophysical methods.

This high throughput, however, comes at a cost: *T*_*m*_ values obtained via TPP reflect a combined effect of protein unfolding and precipitation within the intracellular environment or cellular lysates (Jarzab et al., 2020). Consequently, TPP provides an environment-dependent stability profile that is fundamentally distinct from the thermodynamic stability measurements derived from biophysical assays on purified, isolated proteins. Despite these methodological differences, the sheer scale of sequence-associated thermostability data provided by TPP enabled the development of data-driven melting temperature predictors.

The release of the Meltome Atlas provided an unprecedented volume of sequence-associated *T*_*m*_ data, making it the dominant resource for training deep learning thermostability predictors and shifting the field toward models capable of spanning entire proteomes rather than curated protein families (Dallago et al., 2021). Previous efforts primarily leveraged optimal growth temperatures reported in protein sequence databases as a proxy for melting temperatures. More recent predictors, such as TemStaPro (Pudžiuvelytė et al., 2023), DeepTm (Li et al., 2023), TemBERTure (Rodella et al., 2024), DeepSTABp (Jung et al., 2023), and SaProt (Su et al., 2023), each build on this foundation by coupling Meltome-scale training with protein language model architectures, representing the current state of the art in sequence-based *T*_*m*_ prediction. TemBERTure is built on the protein language model (PLM) ProtBERT-BFD (Elnaggar et al., 2022) and fine-tuned through an adapter-based approach by inserting compact bottleneck layers between transformer layers. It was trained on the Meltome Atlas using 80:10:10 data splits with a 50% sequence identity threshold to reduce potential data leakage. The final model achieved a Mean Absolute Error (MAE) of 6.31 and a coefficient of determination (*R*^2^) of 0.78 on the test set.

DeepSTABp, developed using the same Meltome Atlas, integrates three types of input features through separate neural network blocks: the type of experimental condition (i.e., whether the measurement was conducted in intact cells or lysates), the amino acid sequence embedded by ProTrans-XL, and the optimal growth temperature (OGT) of the organism (Jung et al., 2023). The resulting embeddings from all three blocks are concatenated and passed to a multilayer perceptron (MLP) to predict *T*_*m*_, leading to improved MAE of 3.22 and *R*^2^ of 0.80 on its own test set. However, unlike TemBERTure, DeepSTABp used a less strict random 90:10 train-test split, resulting in higher sequence similarity between training and test sets. Furthermore, the model’s utility is constrained by the requirement for OGT metadata and a rigid dependence on proteomics-specific environmental labels (Cell/Lysate), which limits its generalisability to broader biophysical contexts.

SaProt introduces a structure-aware paradigm by augmenting the amino acid sequence with structure tokens derived from Foldseek (Van Kempen et al., 2024). Each residue is represented as a dual token combining primary sequence information with local 3D geometry (3Di tokens). The model architecture mirrored the 650M-parameter ESM-2 variant and was fine-tuned on the Human-cell split of the Meltome Atlas, achieving a Spearman Correlation Coefficient (SCC) of 0.72. A significant limitation of this model instance is its exclusive training on human proteins, which restricts its predictive range to a narrow window of 40–67°C. Despite this progress, none of these models have been systematically benchmarked across heterogeneous experimental methodologies, leaving their cross-method generalisability largely untested.

In this study, we addressed these limitations by assembling a comprehensive benchmark spanning both proteomics- and biophysics-based *T*_*m*_ datasets, systematically evaluating the generalisability of state-of-the-art (SOTA) deep learning predictors. In addition to the complex predictors mentioned above, we explored simple LoRA-adapted ESM-2 and ESM-3 structural embeddings as competitive approaches to protein *T*_*m*_ prediction. Furthermore, we assessed classification performance as a robust alternative to absolute *T*_*m*_ regression, relevant for the cases in which one seeks to effectively enrich thermostable candidates in enzyme discovery campaigns. We deployed the best-performing model as TmProt 1.0 on Hugging Face Spaces and integrated it into EnzymeMiner 2.0.

## 2 Materials and methods

### 2.1 Data overview

Thermostability datasets can broadly be categorised by the experimental methodology used for their generation.

#### Proteomics-based datasets

To assemble our training dataset, we combined data from two different sources: Meltome Atlas (Jarzab et al., 2020), accessed in June 2023, and ProThermDB (Nikam et al., 2021), accessed in June 2023. The former is a comprehensive dataset containing *T*_*m*_ values for ~48 000 proteins across 13 different organisms: *Thermus thermophilus, Picrophilus torridus, Geobacillus stearothermophilus, Mus musculus, Escherichia coli, Homo sapiens, Bacillus subtilis, Saccharomyces cerevisiae, Drosophila melanogaster, Danio rerio, Arabidopsis thaliana, Caenorhabditis elegans, Oleispira antarctica*. The latter is a database of protein thermostability where organism-based *T*_*m*_ values obtained from LiP-MS experiments (Leuenberger et al., 2017) are deposited along with TPP-based measurements. This data set is less extensive, with only six organisms: *Thermus thermophilus, Escherichia coli, Homo sapiens, Saccharomyces cerevisiae, Arabidopsis thaliana, Toxoplasma gondii*. Protein sequences and their corresponding *T*_*m*_ values from both sources were merged and filtered by length, retaining only sequences between 20 and 2000 amino acids. Duplicate entries were resolved by matching UniProt IDs, retaining the Meltome Atlas record as the primary source. The resulting dataset contains 41 916 proteins from Meltome Atlas and 3525 proteins from ProThermDB. This merged dataset, named **ProMelt**, comprises a total of 45 441 proteins with corresponding *T*_*m*_ values and was used for model training.

#### Biophysics-based datasets

Several datasets were additionally assembled from the databases and literature based on low-throughput biophysical experiments.

BRENDA (Chang et al., 2021) is an enzyme database containing a wide range of functional and biochemical information, including *T*_*m*_ values. A key strength of BRENDA is its manual curation and deposition of individual data points, making it a high-quality resource despite containing fewer *T*_*m*_ records than large-scale datasets such as the Meltome Atlas. We downloaded BRENDA in January 2025 and applied the following filters: only wild-type (WT) proteins were retained, measurements were restricted to neutral pH, and entries annotated with additional experimental conditions, such as the presence of exogenous ligands, ions, or explicit apoenzyme annotations, were excluded to ensure consistency across measurements. The resulting BRENDA subset comprises 321 proteins.

FireProtDB (Stourac et al., 2021) is a manually curated thermostability database that accumulates information on single-point mutations, including changes in Gibbs free energy (ΔΔ*G*) and melting points (Δ*T*_*m*_) between the WT and the mutant protein. Importantly, it also contains *T*_*m*_ values for WT proteins, providing an additional source of high-quality stability data, both literature-derived and experimentally validated entries. We downloaded FireProtDB in June 2023 and obtained a final set of 94 proteins, hereafter referred to as FireProt.

The ERED dataset contains 83 ene reductases with sequence lengths ranging from 390 to 417 residues, encompassing both WT proteins (39 samples) and 44 variants designed via ancestral sequence reconstruction (ASR) to enhance thermostability (Livada et al., 2023). *T*_*m*_ values ranged from 37–55 °C for WT proteins and 39–67 °C for ASR variants. Measurements were conducted using DSF.

The CAS dataset comprises 29 CRISPR-Cas Class II effector proteins (C2EPs) used in the TemStaPro thermostability classifier (Pudžiuvelytė et al., 2023). The *T*_*m*_ values span a range from 33 °C to 67 °C, with protein lengths between 1049 and 1535 residues. Thermostability was assessed using nano differential scanning fluorimetry (nanoDSF).

The HLD dataset consists of 24 haloalkane dehalogenases obtained through genome mining strategies (Vasina et al., 2022). The *T*_*m*_ values range from 35.2 °C to 58.7 °C, with sequence lengths between 271 and 336 residues. Measurements were performed using DSF.

#### Training dataset composition

Proteins shared between ProMelt and the independent test sets, identified by UniProt ID, were removed from ProMelt before training to ensure strict separation between training and evaluation data. The resulting ProMelt dataset (45 441 proteins) was then split into training (77.1%, *n* = 35 054), validation (8.6%, *n* = 3895), and test sets (14.3%, *n* = 6492) at 25% maximum sequence identity using USEARCH (Edgar, 2010) to prevent data leakage. The test set was held out completely from training and validation. To analyse prediction errors against protein length, we sampled 9000 proteins from the training set and 4000 from the test set, stratified to include proteins with both low and high Mean Absolute Error (MAE), using a 5 °C threshold to define the boundary between well-predicted and poorly-predicted examples.

### 2.2 Model training

We explored three simple strategies to predict *T*_*m*_ from protein sequence: (i) the baseline MLP model using protein embeddings from ESM-2 as input, (ii) the ESM-2 model fine-tuned using Low-Rank Adaptation (LoRA), and (iii) the structure-aware modification of the baseline model using ESM-3. The details of the models are provided below.

#### Baseline MLP model

The Baseline MLP model consisted of two hidden layers with 64 and 32 neurons, respectively, with the Rectified Linear Unit (ReLU) activation function. Input features were 1280-dimensional embeddings from ESM-2 (650M) for each protein sequence. The MLP architecture was implemented using pre-built modules from the scikit-learn library (Buitinck et al., 2013).

#### ESM-2 and Low-rank adaptation (ESM2-LoRA)

The protein language model ESM-2 (Lin et al., 2022) (650M) was fine-tuned using Low-Rank Adaptation (Hu et al., 2021) to optimise the inner transformer blocks efficiently by freezing the original ESM-2 weights and introducing trainable low-rank decomposition matrices into the query, key, and value projection layers (Vaswani et al., 2017). Fine-tuning was performed on the ProMelt dataset for a single epoch. We utilised the pre-trained ESM-2 tokeniser with dynamic padding and a batch size of 4. Hyperparameters were optimised using Optuna (Akiba et al., 2019) with the Tree-structured Parzen Estimator sampler over 50 trials, minimising validation RMSE. The following hyperparameters and their search ranges were explored: learning rate (10^−5^–10^−3^, log scale), LoRA rank *r* (*{*1, 2, 4*}*), LoRA alpha (*{*1, 2, 4*}*), dropout (0.0–0.3), weight decay (10^−6^–2 *×* 10^−1^, log scale), gradient clipping (0.0–1.0), and learning rate scheduler (`cosine, linear, constant`). The best configuration used a rank of 1, alpha scaling of 1, dropout of 0.28, weight decay of 1.56 *×* 10^−5^, gradient clipping at 0.805, a cosine learning rate schedule, and an initial learning rate of 4.92 *×* 10^−4^. Training was conducted on an NVIDIA A100 GPU using mixed precision (bf16), and the objective function minimised the mean squared error (MSE) between predicted and experimental *T*_*m*_ values. Model training completed in approximately 70 minutes on the A100 GPU.

#### ESM-3 and Multi-layer Perceptron (ESM3-MLP)

To capture structural information more efficiently, the ESM-3 model (Hayes et al., 2025) (98B) was also used to generate 6144-dimensional structural embeddings per protein. To this end, protein structures were retrieved from AlphaFoldDB (Varadi et al., 2022) where available (accessed January 2025), providing 87.3 % coverage for ProMelt (39 648 proteins). Independent test sets (BRENDA, FireProt, ERED, HLD) predominantly required de novo predictions using OmegaFold (Wu et al., 2022) due to limited AlphaFoldDB availability. The CAS dataset was excluded from the structural embedding pipeline as all its proteins exceed the 1024-residue context window enforced by the ESM-3 structural encoder, with sequence lengths ranging from 1049 to 1535 residues. After generating the embeddings, ESM3-MLP proceeded similarly to the Baseline MLP architecture (two hidden layers: 64/32 ReLU neurons). An overview of the described strategies is shown in Figure 1.

**Fig. 1.**
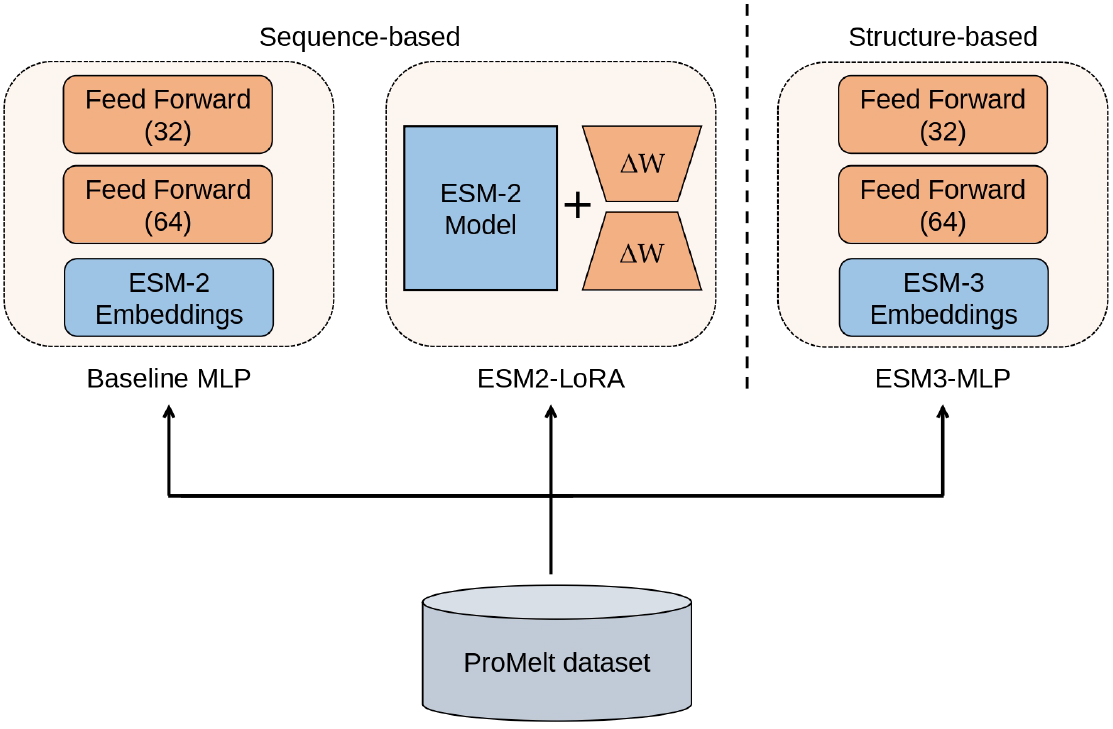
Workflow used for the development of TmProt. The training dataset, ProMelt (comprising 45 441 proteins), was assembled by merging and length-filtering two proteomics-based sources: Meltome Atlas (*n* = 41 916) and ProThermDB (*n* = 3525). The resulting dataset was split at 25% maximum sequence identity into training, validation, and test subsets. Three modelling strategies of increasing complexity were evaluated: Baseline Multi-Layer Perceptron (MLP) uses frozen ESM-2 embeddings as input features to a Baseline MLP regressor with two hidden layers of 64 and 32 neurons, respectively; ESM2-LoRA applies Low-Rank Adaptation (LoRA) fine-tuning directly to ESM-2 transformer blocks for end-to-end thermostability regression, where Δ*W* denotes the trainable low-rank weight updates while the original ESM-2 weights remain frozen; and ESM3-MLP generates structural embeddings from ESM-3 using protein structures retrieved from AlphaFoldDB (Varadi et al., 2022) as input to an MLP regressor with the same architecture as Baseline MLP. The numbers in parentheses indicate the number of neurons in each feed-forward layer. The currently deployed version of TmProt 1.0 is based on strategy ESM2-LoRA (middle).

### 2.3 Model evaluation

#### Evaluation metrics

Four metrics were used to evaluate the performance of the *T*_*m*_ predictor: root mean squared error (RMSE), coefficient of determination *R*^2^, and Pearson’s (PCC) and Spearman’s (SCC) correlation coefficients. In addition to evaluating exact *T*_*m*_ value predictions, we assessed the ability to prioritise thermostable proteins focusing on classification capability, which is critical in enzyme discovery pipelines where identifying highly stable proteins is the main objective. For the combined independent test set (comprising BRENDA, FireProt, ERED, CAS, and HLD), the actual *T*_*m*_ values were binarised using a default threshold of 60 °C: proteins with *T*_*m*_ *≥* 60 °C were considered thermostable (positive), while those below 60 °C were considered non-thermostable (negative). Predicted *T*_*m*_ values were retained as continuous scores to enable threshold-independent evaluation. Model performance was then evaluated using the area under the receiver operating characteristic curve (AUC ROC), the average precision (AP) based on the precision–recall (PR) curve, and the F1 score (harmonic mean of the precision and recall). To quantify the magnitude of the difference between the predicted *T*_*m*_ distributions of thermostable and non-thermostable proteins, we also calculated Cohen’s d as an effect size metric: *d* = (*µ*_1_ *− µ*_2_)*/s*_*pooled*_ where *µ*_1_ and *µ*_2_ are the means of the predicted *T*_*m*_ values for the positive and negative ground-truth classes, respectively, and *s*_*pooled*_ is the pooled standard deviation. This metric provides a scale-independent measure of the model’s ability to separate the two classes, with values above 0.8 generally indicating a large effect size.

#### Thermostability-based top ranking

In a practical enzyme-discovery campaign, a model’s utility is often defined by its ability to enrich a small subset of candidates for high thermostability. To simulate this screening process, we evaluated model performance using a sliding cut-off analysis as follows. For a range of predicted *T*_*m*_ cut-offs, we excluded all proteins predicted to have *T*_*m*_’s below it. From the remaining subset, we calculated the percentage of proteins with an actual experimental *T*_*m*_ *≥* 60 °C. This metric, effectively the enrichment precision, represents the expected success rate when experimentally validating the top-ranked candidates.

#### Structural confidence analysis

To assess whether AlphaFold2 structural reliability correlates with prediction error, we computed per-protein average predicted local distance difference test (pLDDT) scores from AlphaFold2 predictions, excluding terminal residues (first and last 10 positions) due to their inherently lower confidence. These scores were compared against per-protein MAE on the ProMelt test set.

#### Benchmarking against SOTA tools

Three state-of-the-art (SOTA) *T*_*m*_ predictors were selected for benchmarking: TemBERTure, DeepSTABp, and SaProt. TemBERTure was run locally using the official repository^1^, using the TemBERTure regression model (replica 1) with a batch size of 16. DeepSTABp was accessed via its web server^2^ using default parameters: the OGT was left at the server default of 22 °C and the experimental condition was set to Lysate. SaProt predictions were obtained using the Thermostability Model 650M checkpoint (`SaProtHub/Model-Thermostability-650M`), a LoRA adapter fine-tuned on the Human-cell split of the Meltome Atlas, applied on top of the SaProt 650M base model (`westlake-repl/SaProt_650M_AF2`). Since SaProt requires structure-aware sequence tokens, protein structures for each independent test set were first processed using Foldseek to generate combined sequence–structure tokens (3Di), which were then used as input for inference. Unlike the ESM3-MLP pipeline, SaProt imposes no context window restriction on protein length and was therefore evaluated on all five independent test sets, including CAS. Predicted *T*_*m*_ values were reverse-normalised from the FLIP (Dallago et al., 2021) normalisation range of 40–67 °C to recover absolute temperatures. All three tools were evaluated using identical evaluation metrics.

## 3 Results

### 3.1 Preliminary data analysis

We first examined the *T*_*m*_ distributions of the five independent datasets (Figure 2). Among them, BRENDA and FireProt datasets contained a substantial fraction of stable proteins (*T*_*m*_ > 60 °C).

**Fig. 2.**
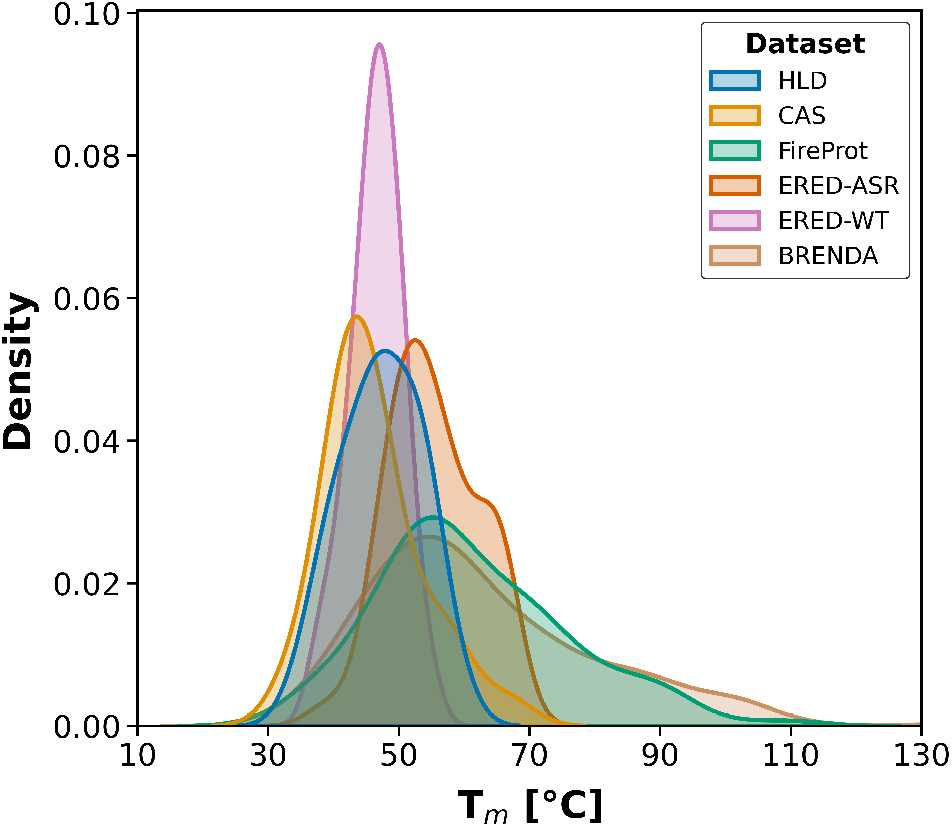
Melting temperature (*T*_*m*_) distribution of the biophysics-based independent datasets. Notably, the ancestral sequence reconstruction (ERED-ASR, orange) variants exhibit a clear shift toward higher temperatures compared to the wild-type (ERED-WT, pink) enzymes, reflecting their enhanced thermostability.

To assess potential measurement discrepancies across the experimental technique, we compared the *T*_*m*_ values of the 60 overlapping proteins shared between the ProMelt and BRENDA datasets (Figure 3a). These proteins had the same UniProtID but were measured by different experimental techniques: TPP and LiP-MS for the ProMelt, and biophysical for BRENDA. We observed limited agreement between the two datasets, with a moderate positive correlation (*r* = 0.57) but substantial deviations in the reported *T*_*m*_ values (Figure 3a). A similar comparison of the 29 shared proteins in the ProMelt and FireProt datasets revealed virtually no agreement (*r* = 0.05) and large differences in measured *T*_*m*_ values (Figure 3a). In contrast, the biophysics-based datasets showed strong agreement with each other (Figure 3b). To ensure strict separation between training and evaluation data, all proteins shared between ProMelt and the independent test sets (BRENDA, FireProt, ERED, CAS, HLD), identified by UniProt ID, were removed from ProMelt before model training. Additionally, the eight proteins shared between FireProt and BRENDA were removed from BRENDA to ensure the two datasets remain fully disjoint for independent evaluation.

**Fig. 3.**
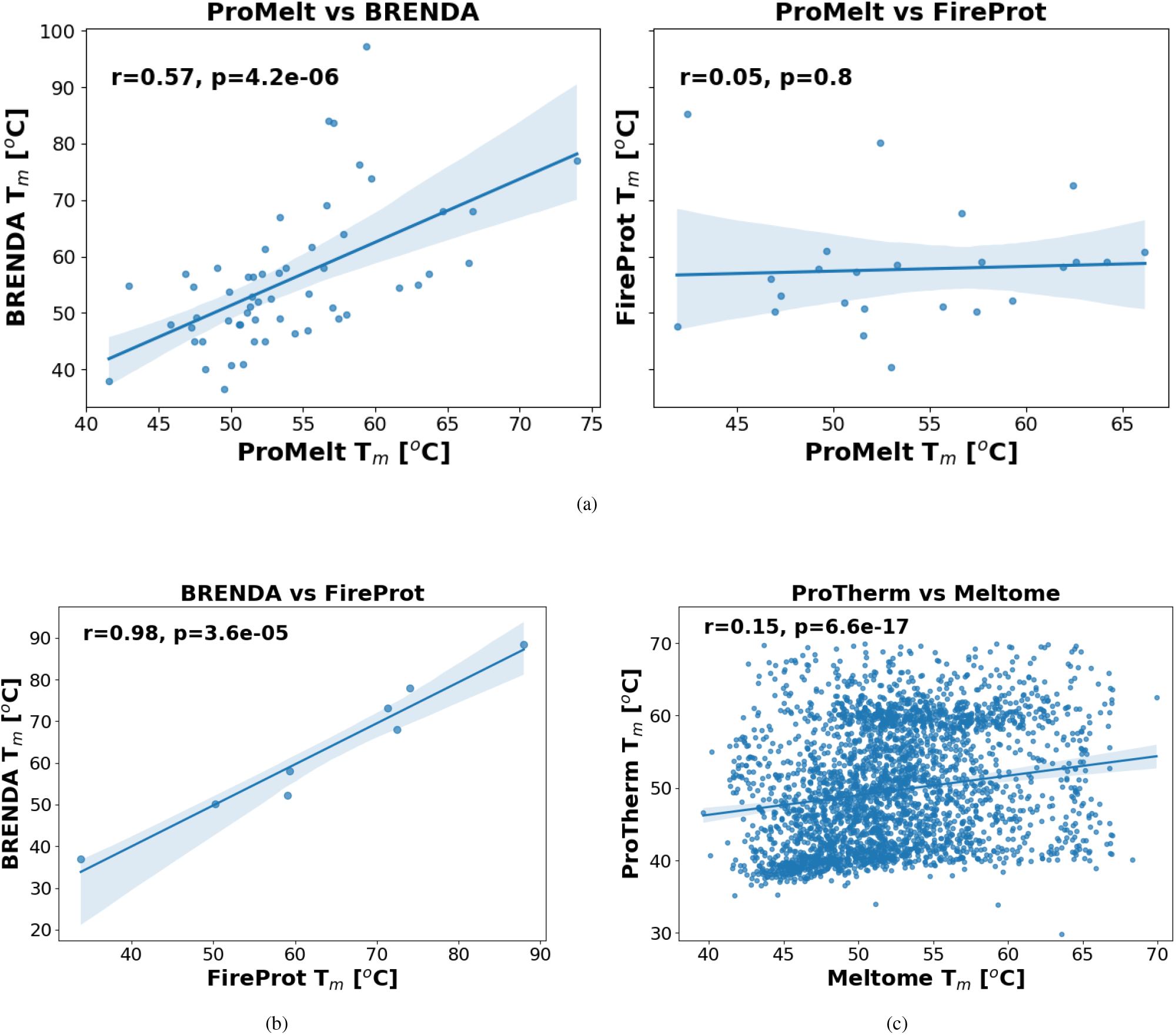
Comparison of *T*_*m*_ values across protein thermal stability datasets. **(a)** Proteomics-based ProMelt shows weak agreement with biophysics-based BRENDA (*r* = 0.57) and FireProt (*r* = 0.05) datasets. **(b)** Strong agreement between biophysics-based BRENDA and FireProt (*r* = 0.98) confirms internal consistency within this measurement class. **(c)** Poor agreement between two proteomics-based methods, Meltome Atlas (TPP) and ProTherm (LiP-MS) (*r* = 0.15), indicates that experimental protocol substantially affects reported *T*_*m*_ values.

Notably, even within the proteomics domain, different approaches yielded inconsistent measurements: the 3140 proteins shared between the Meltome Atlas (TPP) and ProTherm (LiP-MS) showed a weak correlation (*r* = 0.15, Figure 3c), suggesting that proteomics-based *T*_*m*_ measurements are sensitive to the specific experimental protocol employed. To avoid redundancy, overlapping proteins were removed from ProTherm, keeping Meltome Atlas as the primary source.

### 3.2 Melting temperature prediction

Three modelling strategies of increasing complexity were trained and compared using the proteomic-based ProMelt dataset. The Baseline MLP operates on sequence-derived features without pretrained protein representations, serving as a reference for the performance achievable without large-scale pretraining. The ESM2-LoRA model fine-tunes the ESM2 protein language model using LoRA, allowing the pretrained sequence representations to be updated efficiently on the thermostability task while avoiding full-model fine-tuning. The ESM3-MLP model keeps the ESM3 encoder frozen and trains a lightweight MLP regression head on top of its representations, which encode not only sequence but also predicted structural context, allowing us to assess whether structural information embedded in ESM3 provides additional predictive signal beyond sequence alone.

The Baseline MLP model evaluated on the ProMelt test set achieved moderate performance across all metrics (Table 1). The ESM2-LoRA model showed improved ranking ability over the baseline, reflected in higher *R*^2^, PCC, and SCC, despite a slightly higher RMSE. The ESM3-MLP model achieved the best overall performance on the ProMelt test set across all four metrics (Table 1), confirming that structural context contributes meaningfully to predicted thermostability. While the direct benchmarking against the state-of-the-art tools is complicated due to the lack of established data splits and metrics, our values in Table 1 are in line with those reported by other methods, e.g., the PCC of 0.78-0.90 (Elnaggar et al., 2022; Jung et al., 2023).

**Table 1.** Performance of the three proposed models on the ProMelt test set.

| Model | RMSE | $R^2$ | PCC | SCC |
| --- | --- | --- | --- | --- |
| Baseline MLP | 5.78 | 0.67 | 0.82 | 0.59 |
| ESM2-LoRA | 6.23 | 0.69 | 0.83 | 0.62 |
| ESM3-MLP | **5.40** | **0.73** | **0.85** | **0.66** |

RMSE, root mean squared error; $R^2$ , coefficient of determination. PCC, Pearson correlation coefficient; SCC, Spearman's rank correlation coefficient.

#### Analysis of prediction errors

To identify specific factors influencing prediction error in *T*_*m*_estimation, we conducted a systematic error analysis using the Baseline MLP model as a controlled reference on the ProMelt test set. We examined the effects of sequence clustering (USEARCH cluster size), protein length, organism origin, embedding space structure (via t-SNE), and structural confidence (AlphaFold2 pLDDT scores). None of these factors showed a consistent association with prediction error (Figures S2– 4; cluster size *r* = *−*0.03, protein length *r* = *−*0.06). As shown in Figure 4, only a weak positive correlation between pLDDT and MAE was observed (*r* = 0.26), indicating that structural confidence shows minimal association with prediction error. Taken together, these analyses suggest that the broad error distribution is not attributable to any single identifiable factor, pointing instead to inherent noise in the TPP-derived *T*_*m*_ labels as the most likely source of irreducible error.

**Fig. 4.**
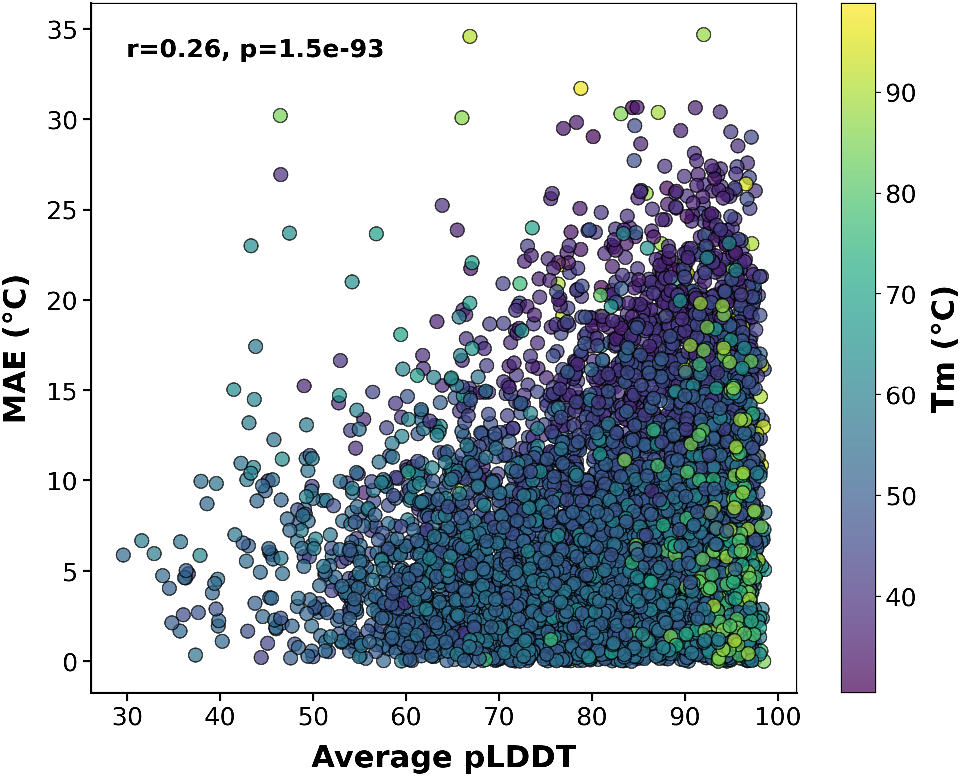
Mean Absolute Error (MAE) per cluster vs. average Predicted Local Distance Difference Test (pLDDT) confidence score for proteins in the ProMelt test set, coloured by *T*_*m*_.

#### Evaluation on the biophysical datasets

Despite the promising performance on the ProMelt test dataset, our evaluation of the model on five external biophysical datasets showed a substantial drop in performance (e.g., ESM3-MLP *R*^2^ fell from 0.73 on ProMelt to 0.35 on BRENDA and 0.23 on FireProt). On the larger BRENDA and FireProt datasets, both ESM2-LoRA and ESM3-MLP outperformed the Baseline MLP and achieved SCC of 0.43–0.56 (Table 2). Performance deteriorated markedly on the ERED and HLD datasets, which represent single enzyme families (ene reductases and haloalkane dehalogenases, respectively) containing evolutionarily related sequences within each family. Here, all models produced negative *R*^2^ values, reflecting the inherent difficulty of predicting fine-grained stability differences among closely homologous variants.

**Table 2.**
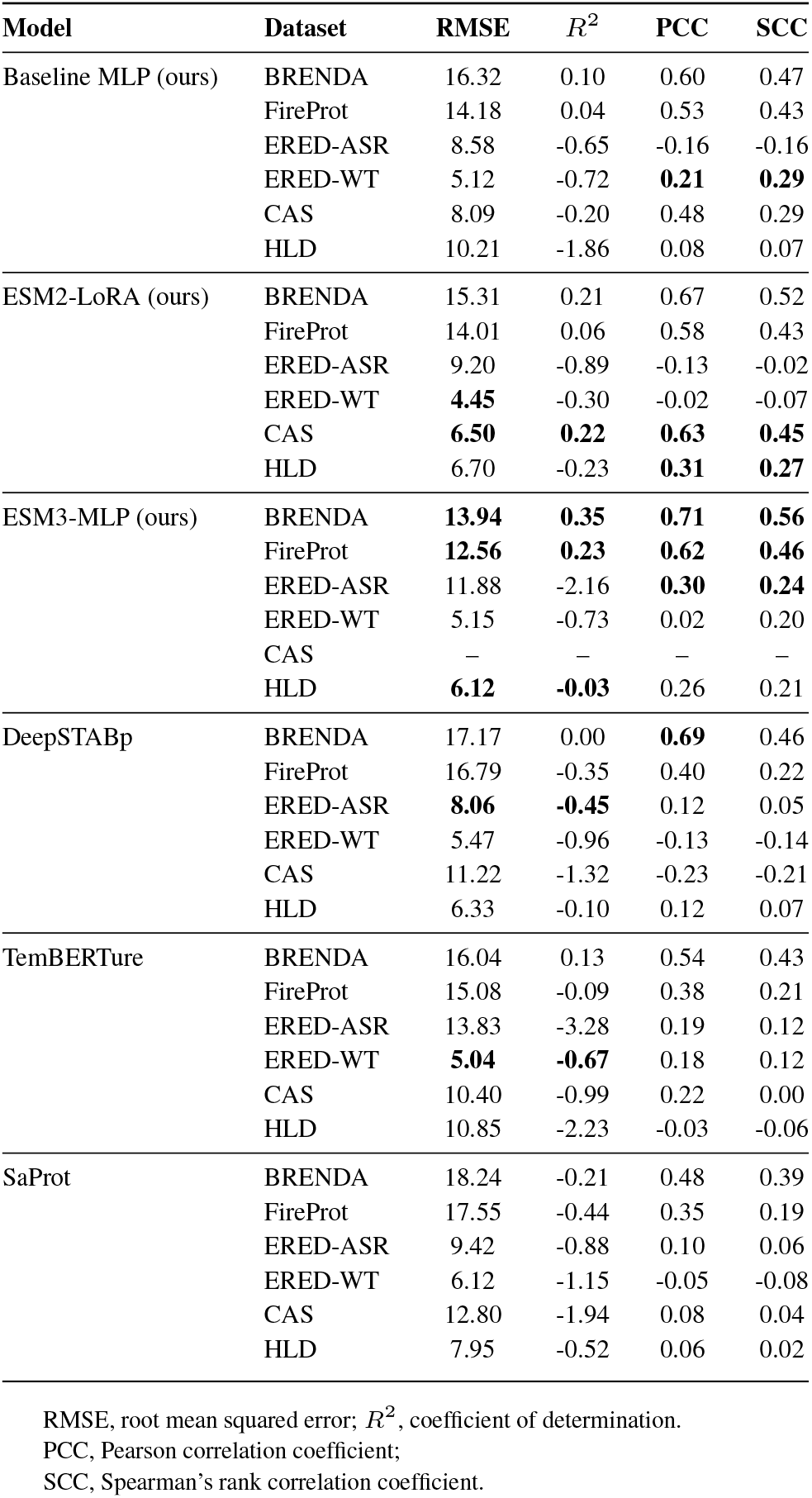
Performance comparison of all models across independent biophysics test sets. ESM3-MLP metrics for the CAS dataset are omitted due to the structural context window limitations (see Section 2.2).

We further evaluated the state-of-the-art *T*_*m*_ predictors, DeepSTABp, TemBERTure, and SaProt, across multiple external datasets. Table 2 summarises the comparative evaluation results in terms of RMSE, *R*^2^, PCC, and SCC.

Apart from a few metrics, all three predictors showed lower performance compared to our ESM2-LoRA and ESM3-MLP models. DeepSTABp exhibited moderate performance on the BRENDA dataset but showed substantially higher errors and lower correlation metrics on FireProt, CAS, and HLD datasets. TemBERTure generally performed better than DeepSTABp on BRENDA and FireProt but worse on ERED, CAS, and HLD datasets, indicating variable generalisability across different data sources. Despite incorporating structural information, SaProt did not surpass the other tools, possibly due to its training focus on human-specific proteins. By contrast, ESM2-LoRA and ESM3-MLP yielded higher regression accuracy and stronger correlation coefficients than all three SOTA predictors across most benchmarks, showing that parameter-efficient fine-tuning of ESM embeddings can match or exceed more complex predictors on cross-method benchmarks.

### 3.3 Thermostable Protein Identification

Given the difficulty of predicting absolute *T*_*m*_ values for biophysics-based datasets, we further investigated whether models can perform well in protein screening campaigns, which usually require efficient protein prioritisation rather than exact *T*_*m*_ value prediction. To this end, we evaluated the ability of all models to distinguish thermostable proteins across the combined independent biophysics datasets using a cut-off of *T*_*m*_ *≥* 60 °C, a commonly adopted benchmark in industrial biotechnology where enzymes must remain functional at elevated temperatures (Rigoldi et al., 2018). Classification performance is summarised in Figure 5 and Table 3.

**Table 3.** Classification performance and effect sizes for thermostability identification on the combined independent biophysics datasets (cut-off = 60 °C). Cohen’s *d* indicates the magnitude of separation between predicted distributions of thermostable and non-thermostable proteins. *ESM3-MLP excludes the CAS dataset due to the structural context window limitation of the ESM-3 encoder (see Section 2.2).

| Model | AUC | AP | F1 | Cohen’s $d$ |
| --- | --- | --- | --- | --- |
| Baseline MLP | 0.72 | 0.68 | 0.61 | 0.99 |
| ESM2-LoRA | 0.75 | 0.69 | 0.61 | 1.08 |
| ESM3-MLP* | **0.77** | **0.74** | **0.65** | **1.17** |
| SaProt | 0.65 | 0.51 | 0.57 | 0.51 |
| DeepSTABp | 0.66 | 0.65 | 0.56 | 0.90 |
| TemBERTure | 0.72 | 0.65 | 0.60 | 0.92 |

AUC, area under the receiver operating characteristic curve; AP, average precision; F1, harmonic mean of precision and recall; Cohen’s $d$ , standardised effect size measuring separation between predicted $T_m$ distributions of thermostable and non-thermostable proteins.

**Fig. 5.**
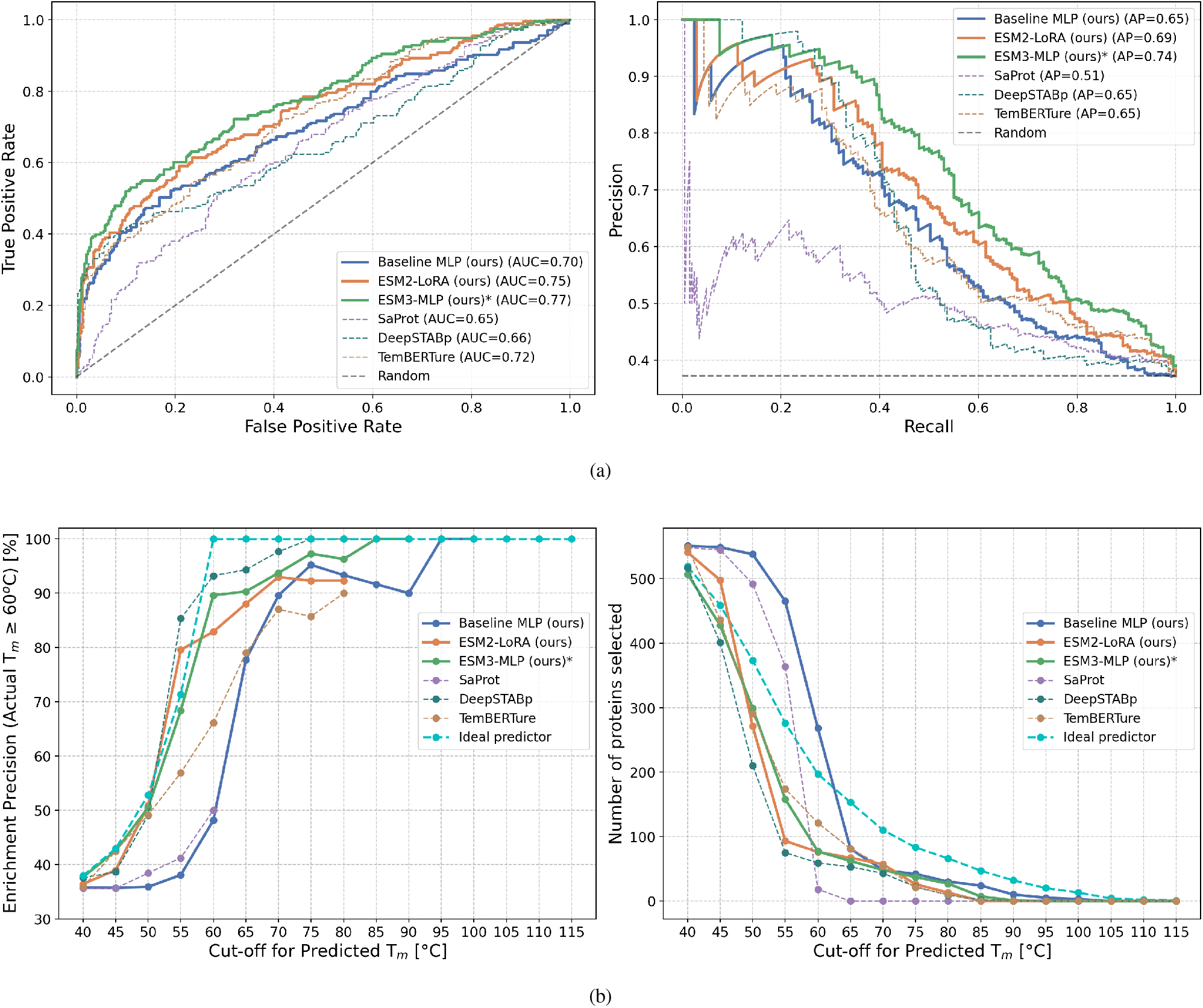
Classification performance for thermostable protein identification on the combined independent biophysics datasets (*T*_*m*_ *≥* 60 °C). **(a)** ROC curve (left) and precision–recall curve (right); ESM3-MLP achieves the highest discrimination (AUC = 0.77, AP = 0.74). **(b)** Enrichment precision (left) and number of retained proteins (right) as a function of the predicted *T*_*m*_ cut-off; the cyan dashed line indicates ideal predictor performance. *ESM3-MLP excludes the CAS dataset due to the structural context window limitation of the ESM-3 encoder. ROC: receiver operating characteristic; AUC: area under the ROC curve; AP: average precision.

Our models consistently outperformed all three SOTA predictors (DeepSTABp, TemBERTure, SaProt) across all classification metrics. ESM3-MLP achieved the highest discrimination performance (AUC = 0.77, AP = 0.74, Cohen’s d = 1.17), followed by ESM2-LoRA (AUC = 0.75, AP = 0.69, Cohen’s d = 1.08) and the Baseline MLP (AUC = 0.72, AP = 0.68, Cohen’s d = 0.99). The large Cohen’s d values (*≥*0.8) for all three of our models indicate strong separation between predicted *T*_*m*_ distributions of thermostable and non-thermostable proteins, enabling reliable identification of thermostable candidates. DeepSTABp and TemBERTure showed moderate performance, though consistently below our approaches (Table 3).

Analysis of the enrichment precision curves (Figure 5b) confirms that filtering candidate proteins by predicted *T*_*m*_ is a practically useful strategy: as the predicted *T*_*m*_ cut-off increases, the fraction of truly thermostable proteins in the retained subset rises steadily for all three of our models, reaching near-complete enrichment above 75 °C. Thus, although absolute *T*_*m*_ regression remains sensitive to cross-method discrepancies, our models retain sufficient discriminatory power to reliably enrich candidate sets for downstream experimental validation. Notably, DeepSTABp reaches full enrichment at lower predicted thresholds than our models, likely reflecting a systematic underestimation of absolute *T*_*m*_ values: proteins predicted at 60 °C by DeepSTABp may correspond to truly thermostable proteins that our models assign higher scores to.

#### TmProt 1.0 deployment

To facilitate the access to *T*_*m*_ prediction, we also implemented TmProt 1.0 as a web tool. We selected to deploy ESM2-LoRA (not ESM3-MLP) as TmProt 1.0 due to its higher inference speed, no input sequence length limits (ESM3 is constrained to 1024 residues), and robustness to sequence-only input, which is common in screening pipelines. The web server accepts batch FASTA submission and returns predicted *T*_*m*_ and binary classification (*T*_*m*_ *≥* 60 °C) for each sequence. It is hosted at https://loschmidt.chemi.muni.cz/tmprot/ and integrated into EnzymeMiner 2.0 (https://loschmidt.chemi.muni.cz/enzymeminer/). All models and training scripts are also available at https://github.com/loschmidt/TmProt.

## 4 Discussion

The central finding of this study concerns measurement methodology rather than model design: *T*_*m*_ values derived from TPP and those obtained from biophysical assays on purified proteins are not interchangeable as prediction targets. The near-zero correlation observed between ProMelt and FireProt measurements for proteins with matched UniProt IDs (*r* = 0.05, Figure 3a) indicates that the two measurement regimes capture partially distinct biological quantities. TPP-derived *T*_*m*_ is an environment-dependent quantity, modulated by interactions with small molecules, metabolites, and binding partners, as well as post-translational modifications such as phosphorylation (Mateus et al., 2020; Jarzab et al., 2020). Biophysical measurements on purified proteins remove the cellular context captured by TPP; however, even under controlled conditions, measured *T*_*m*_ values remain sensitive to buffer composition, pH, and ionic strength, which can vary across individual measurements deposited in the database (Chang et al., 2021). Although some performance loss is expected when transferring between these regimes, the magnitude of the gap is striking: RMSE exceeds 10 °C and *R*^2^ is negative on several datasets. This concern extends beyond the models evaluated here since TemBERTure (Rodella et al., 2024), DeepSTABp (Jung et al., 2023), and SaProt (Su et al., 2023) all rely on the Meltome Atlas (Jarzab et al., 2020) as their primary training source, meaning the cross-method generalization problem is shared across the current generation of sequence-based *T*_*m*_ predictors.

The apparent advantage of our models over the SOTA predictors on the biophysical test sets should not be overestimated. None of the evaluated models, including our own, were trained or fine-tuned on biophysical data. Moreover, DeepSTABp was evaluated at a fixed OGT of 22 °C and Lysate condition, the server defaults that do not reflect the experimental conditions of the biophysical test sets. SaProt was trained exclusively on the human-cell split of the Meltome Atlas (6722 proteins), narrowing down its predictive range. ESM2-LoRA (a simple parameter-efficient fine-tune) matches or exceeds more complex models across benchmarks. The selected LoRA configuration used rank r=1, the smallest tested, suggesting that minimal parameter updates suffice for fine-tuning ESM-2 to this task.

Despite the overall performance gap, ranking proteins by predicted *T*_*m*_ proved more robust than absolute *T*_*m*_ regression in cross-method evaluation. The enrichment precision analysis (Figure 5b) confirms that predicted *T*_*m*_ scores can reliably prioritise thermostable candidates above a classification threshold, the relevant operating mode in enzyme discovery campaigns (Hon et al., 2020).

As far as ProMelt performance is concerned, our error distribution analysis revealed that prediction errors were broadly distributed across protein embedding space, organism origin, sequence identity, and protein length, with no single factor accounting for the high-error cases (Figure S2–S This is yet to be investigated further if such error distributions stem from the irreducible label noise in TPP-derived *T*_*m*_ values or due to systematic deficiencies in the model. Two strategies may elucidate this. First, a self-distillation approach analogous to the second training round in AlphaFold2 (Jumper et al., 2021), in which the model is trained to predict its own errors, could provide per-protein reliability estimates and allow users to identify predictions likely to be unreliable. Second, pre-screening of protein embeddings prior to inference (Prabakaran and Bromberg, 2026), which scores each representation against its nearest neighbours in latent space to quantify biological relevance, could flag proteins likely to produce unreliable predictions before the prediction step.

The independent biophysical test sets used in this study, while carefully curated, are insufficient in scale to serve as training data for biophysics-aware models. The recently released FireProtDB 2.0 (Musil et al., 2026), which aggregates absolute *T*_*m*_ measurements alongside mutational stability data from multiple sources, provides a growing collection of high-quality thermostability measurements that could support fine-tuning in future work. Test-time training has recently been demonstrated to improve generalisation of protein language models on structure and fitness prediction tasks (Bushuiev et al., 2026), applying an analogous self-supervised adaptation strategy to ESM2-LoRA at inference time may similarly improve *T*_*m*_ prediction for proteins from biophysical measurement regimes not seen during training.

## 5 Conclusions

The core limitation of existing *T*_*m*_ predictors is their evaluation on a single training source: the Meltome Atlas, which measures protein stability in cellular lysates via TPP. We assembled five independent biophysics-based datasets (BRENDA, FireProt, ERED, CAS, HLD) measured by DSF and DSC on purified, isolated proteins. This cross-methodology evaluation revealed that *T*_*m*_ values from TPP and biophysical assays are not interchangeable, and no existing model generalises well across both regimes.

Our comparative analysis of ProMelt, BRENDA, and FireProt datasets revealed pronounced discrepancies between *T*_*m*_ values obtained from cellular lysates and those obtained from purified proteins.

Across the evaluated benchmarks, the LoRA-adapted ESM-2 and ESM-3 structural models generally outperformed the Baseline MLP and current state-of-the-art predictors. Although ESM-3 provides maximum accuracy through structural context, ESM2-LoRA achieved comparable robustness with significantly higher computational efficiency. Consequently, we selected ESM2-LoRA for deployment as TmProt 1.0, a user-friendly web server for protein melting temperature prediction.

Despite these improvements, no single model consistently dominated across all datasets. Beyond regression of absolute *T*_*m*_ values, our models demonstrated high discriminatory power for identifying thermostable proteins (*T*_*m*_ *≥* 60 °C). Ranking by predicted *T*_*m*_ with a user-defined cut-off proved a practical strategy for enzyme mining, directly supported by TmProt 1.0, available at https://loschmidt.chemi.muni.cz/tmprot/ and EnzymeMiner 2.0 (https://loschmidt.chemi.muni.cz/enzymeminer/).

## Acknowledgements

Computational resources were provided by the e-INFRA CZ [LM2018140] and ELIXIR-CZ [LM2023055] projects, supported by the Ministry of Education, Youth and Sports of the Czech Republic. Brno Ph.D. Talent Scholarship holder Pavel Kohout acknowledges funding from the Brno City Municipality. This project is supported by the CLARA project (European Union’s Horizon Europe research and innovation programme under Grant Agreement No. 101136607) and by CLARA-OP JAK (No. 02_23_029 Teaming-CZ II). This work reflects only the authors’ view, and the European Commission is not responsible for any use that may be made of the information it contains. We are grateful to Miloš Musil for collecting the FireProt dataset used in this study.

## Supplementary data

The following supplementary figures provide additional characterisation of the ProMelt dataset and a systematic analysis of prediction errors. ProMelt spanned a wide range of *T*_*m*_ values, from 27 °C to 99 °C, and included samples from thermophilic and psychrophilic organisms (see Section 2.1 of the main text and Figure S1). Figures S2–S5 examine candidate sources of prediction error (sequence clustering, protein length, organism origin, and ESM-2 embedding space). Figures S6 and S7 present per-dataset classification performance at the 60 °C thermostability cut-off.

## Data availability

CSV files containing *T*_*m*_ values, UniProt IDs, and protein sequences for the ProMelt training dataset and independent test sets (BRENDA, FireProt, ERED, CAS, HLD), along with corresponding FASTA files, are available on Zenodo at [DOI]. All data processing scripts and curated datasets are provided to ensure full reproducibility of the TmProt framework.

---

**Fig. S1.**
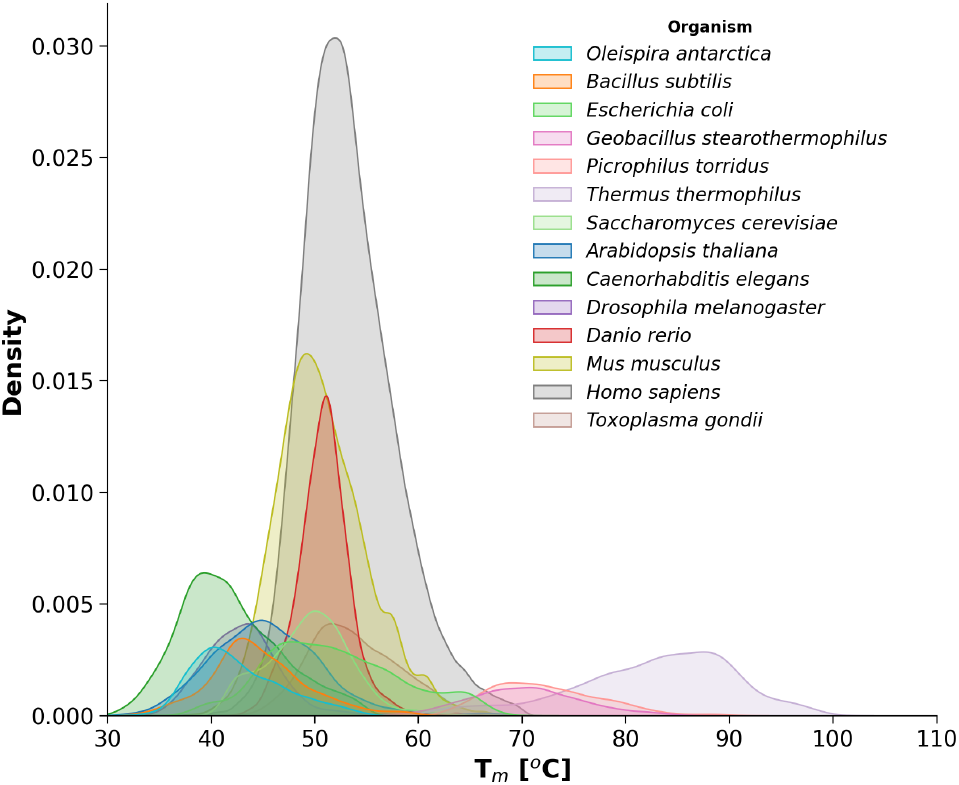
Distribution of *T*_*m*_ values per organism in the ProMelt dataset. Most data points originate from human cell lines.

**Fig. S2.**
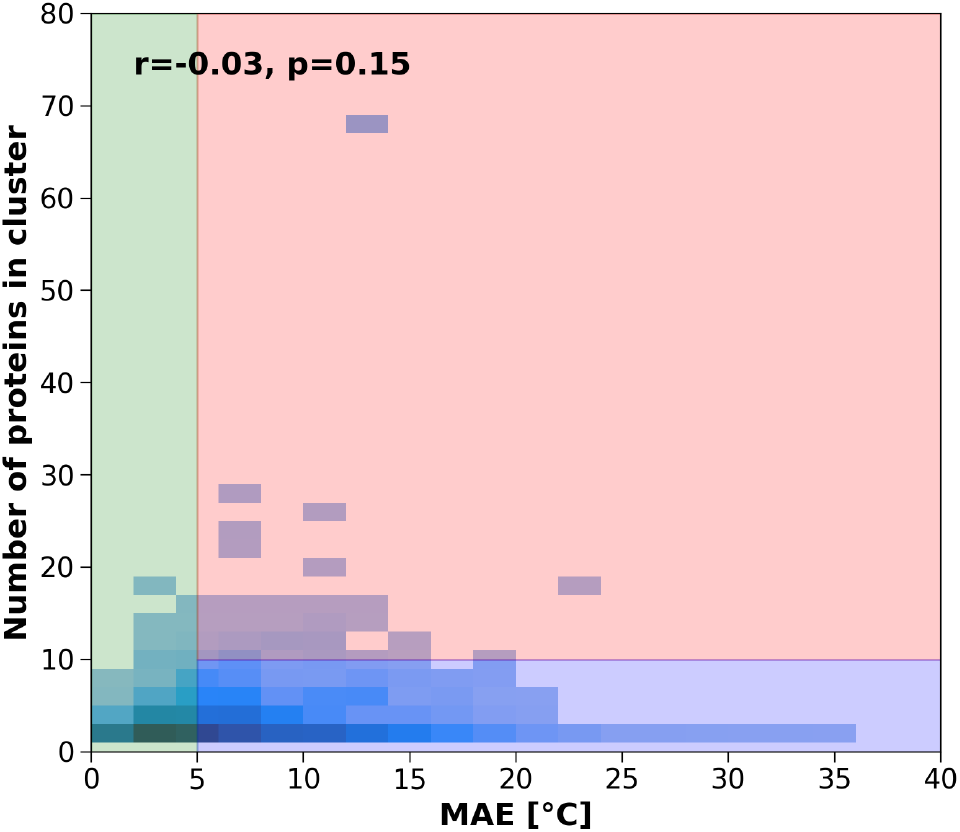
Histogram of Mean Absolute Error (MAE) versus cluster population, visually divided into 3 sectors by colour. Ideally, all the predictions would fall into the green area, corresponding to the model prediction error below 5 °C, regardless of the number of proteins in each cluster. The blue region corresponds to high errors for clusters with few sequences, which might be expected due to a low amount of similar proteins to learn from. The red sector represents the main challenge, as the model failed to recognise patterns for clusters with a sufficient number of similar proteins. The near-zero correlation (*r* = *−*0.03) indicates that cluster size does not explain prediction error, and high-error cases occur regardless of how many similar sequences were available during training.

**Fig. S3.**
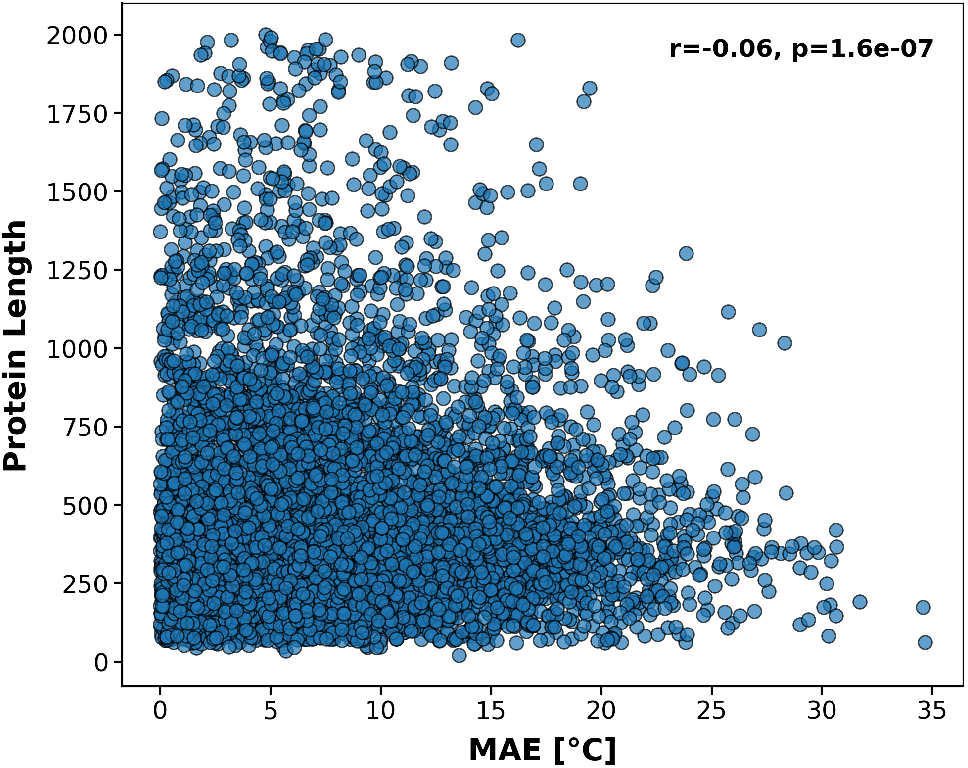
Mean Absolute Error (MAE) against protein length. The near-zero correlation (*r* = *−*0.06) indicates that sequence length does not systematically drive prediction error; proteins of all lengths contribute to the high-error tail, suggesting that length is not a confounding factor in model performance.

**Fig. S4.**
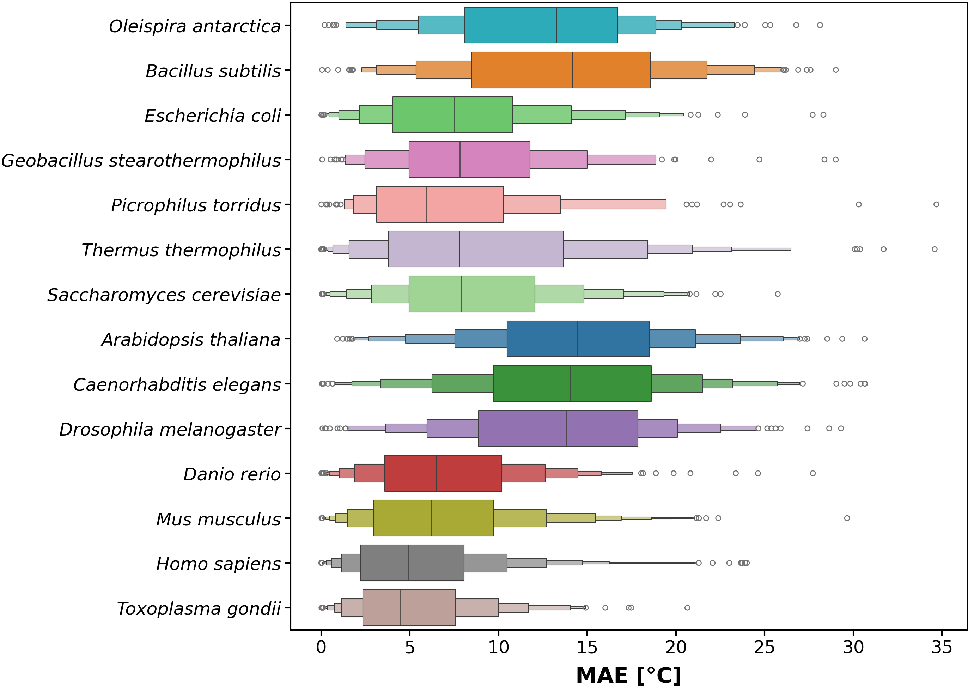
Distribution of Mean Absolute Error (MAE) values per organism. While extremophiles (*T. thermophilus, P. torridus, G. stearothermophilus*) display broad error distributions, absolute error ranges are comparable to those for proteins from mesophilic and psychrophilic organisms, suggesting that organism type alone does not determine prediction accuracy.

**Fig. S5.**
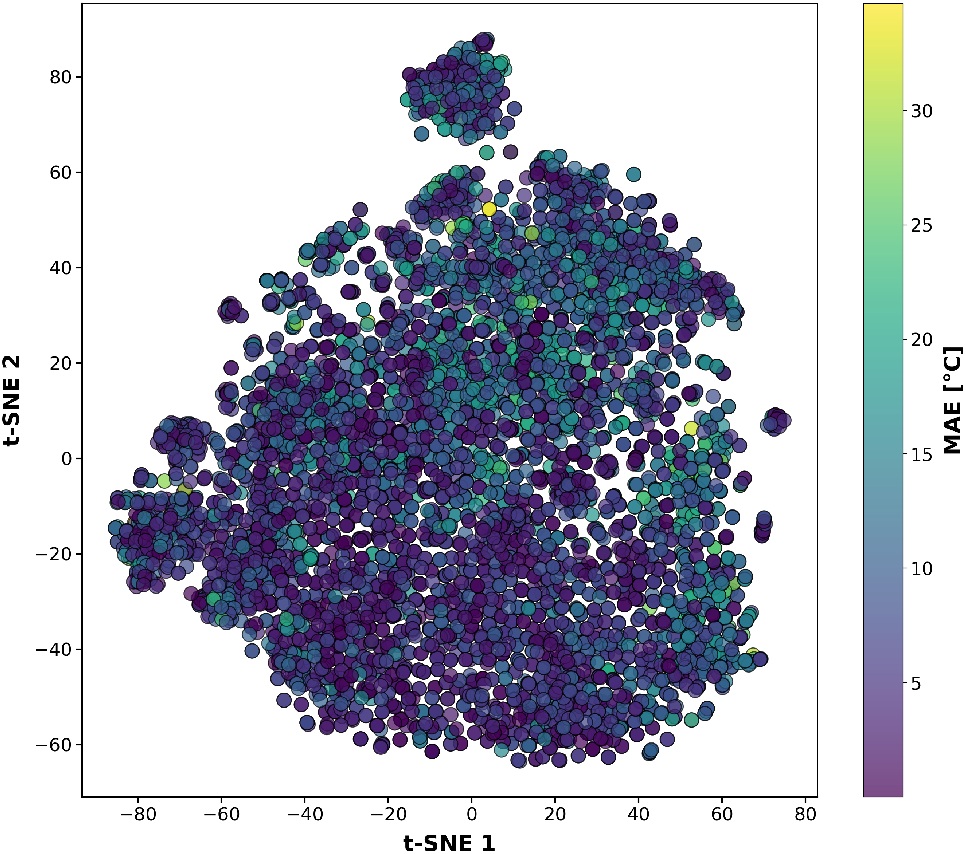
Two-dimensional scatter plot of ESM-2 embeddings projected using t-SNE (van der Maaten and Hinton, 2008). Each point represents a protein from the test (left) and training (right) sets, coloured by prediction MAE. No spatially coherent clusters of high error were observed, indicating that prediction failures are distributed broadly across sequence space rather than being confined to specific protein families.

**Fig. S6.**
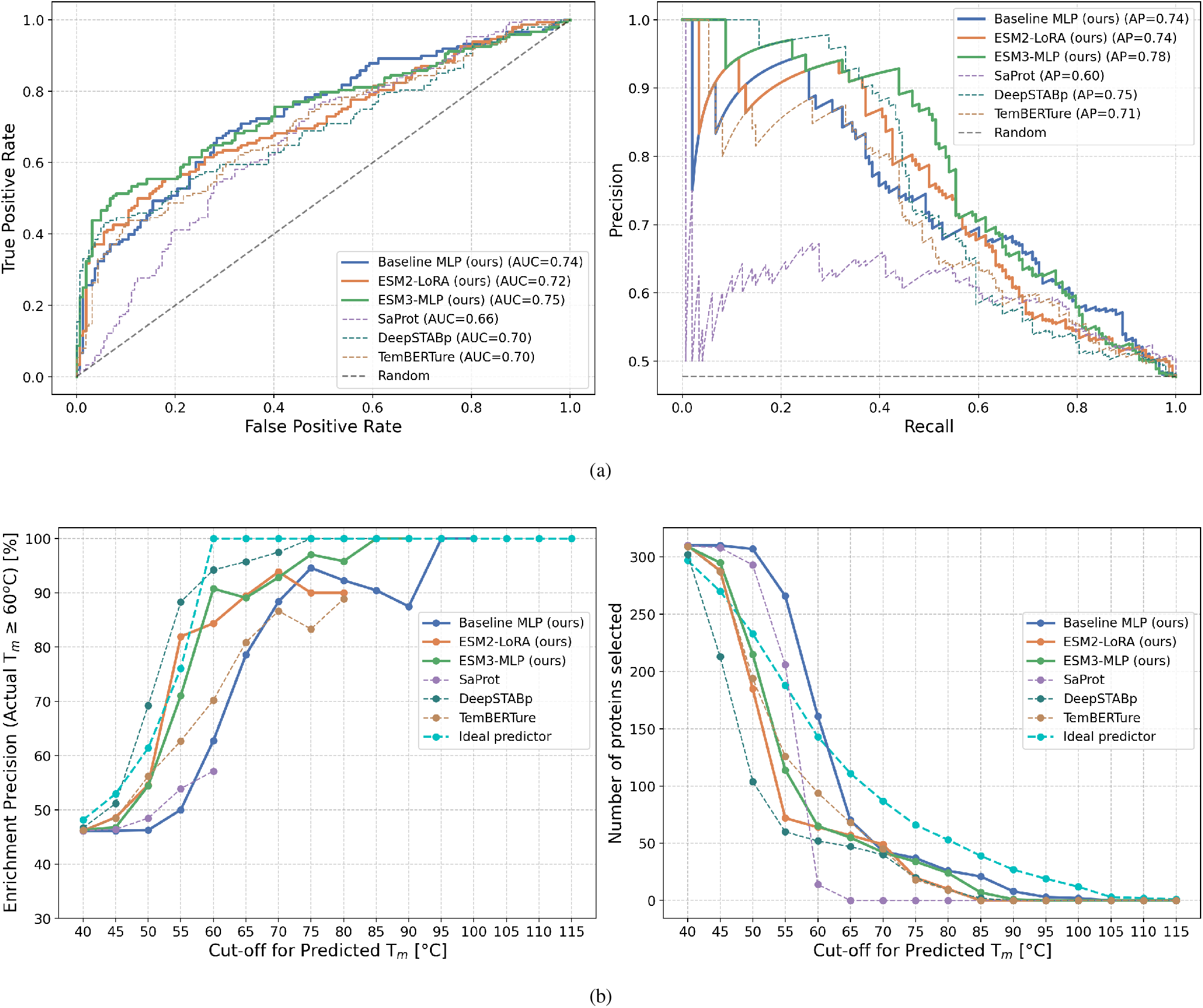
Classification performance on the BRENDA dataset for thermostable protein identification (*T*_*m*_ *≥* 60 °C). **(a)** ROC curve (left) and precision–recall curve (right); ESM3-MLP achieves the highest discrimination (AUC = 0.75, AP = 0.78). **(b)** Enrichment precision (left) and number of retained proteins (right) as a function of the predicted *T*_*m*_ cut-off; the cyan dashed line indicates ideal predictor performance. *ESM3-MLP excludes the CAS dataset due to the structural context window limitation of the ESM-3 encoder (see Section 2.2 of the main text). ROC: receiver operating characteristic; AUC: area under the ROC curve; AP: average precision.

**Fig. S7.**
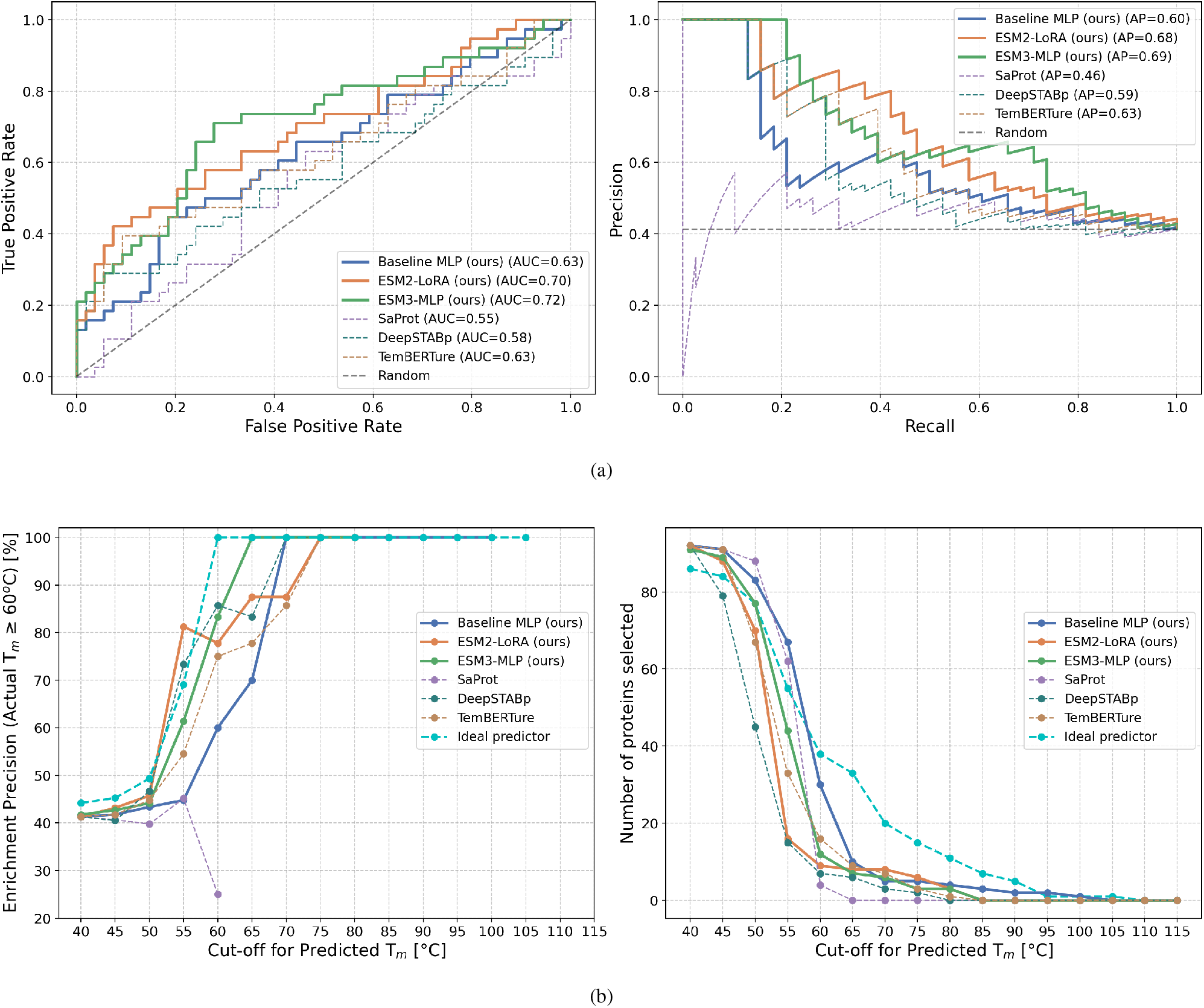
Classification performance on the FireProt dataset for thermostable protein identification (*T*_*m*_ *≥* 60 °C). **(a)** ROC curve (left) and precision–recall curve (right); ESM3-MLP achieves the highest discrimination (AUC = 0.72, AP = 0.69). **(b)** Enrichment precision (left) and number of retained proteins (right) as a function of the predicted *T*_*m*_ cut-off; the cyan dashed line indicates ideal predictor performance. *ESM3-MLP excludes the CAS dataset due to the structural context window limitation of the ESM-3 encoder (see Section 2.2 of the main text). ROC: receiver operating characteristic; AUC: area under the ROC curve; AP: average precision.

## Footnotes

1 https://github.com/ibmm-unibe-ch/TemBERTure

2 https://csb-deepstabp.bio.rptu.de/

